# Global Profiling of the Lysine Crotonylome in Different Pluripotent States

**DOI:** 10.1101/2020.09.21.288167

**Authors:** Yuan Lv, Chen Bu, Jin Meng, Carl Ward, Giacomo Volpe, Jieyi Hu, Mengling Jiang, Lin Guo, Jiekai Chen, Miguel A. Esteban, Xichen Bao, Zhongyi Cheng

## Abstract

Pluripotent stem cells (PSCs) can be expanded *in vitro* in different culture conditions, resulting in a spectrum of cell states with distinct properties. Understanding how PSCs transition from one state to another, ultimately leading to lineage-specific differentiation, is important for developmental biology and regenerative medicine. Although there is significant information regarding gene expression changes controlling these transitions, less is known about post-translational modifications of proteins. Protein crotonylation is a newly discovered post-translational modification where lysine residues are modified with a crotonyl group. Here, we employed affinity purification of crotonylated peptides and liquid chromatography-tandem mass spectrometry (LC-MS/MS) to systematically profile protein crotonylation in mouse PSCs in different states including ground, metastable and primed state, as well as metastable PSCs undergoing early pluripotency exit. We successfully identified 3628 high-confidence sites in 1426 proteins. These crotonylated proteins are enriched for factors involved in functions related to pluripotency such as RNA biogenesis, central carbon metabolism and proteasome function. Moreover, we found that increasing the cellular levels of crotonyl-CoA through crotonic acid treatment promotes proteasome activity in metastable PSCs and delays their differentiation, consistent with previous observations showing that enhanced proteasomal activity helps to sustain pluripotency. Our atlas of protein crotonylation will be valuable for further studies of pluripotency regulation and may also provide insights into the role of metabolism in other cell fate transitions.

## Introduction

Pluripotency is a transient state of the developing embryo. Through this property, cells in the inner cell mass (ICM) of the blastocyst have the capacity to differentiate into all tissues that compose the body [1]. *In vitro*, pluripotency can be maintained by culturing ICM cells in specific conditions, which then produces PSCs. Accordingly, PSCs are self-renewing, can be maintained indefinitely *in vitro* and differentiated on demand upon exposure to specific signalling cues. Studying the regulation of pluripotency maintenance and exit in PSCs is important not only to understand embryonic development but also for cell fate transitions in other contexts (*e.g*., somatic cell reprogramming and transdifferentiation, cancer, or aging). Mouse PSCs are a well-studied pluripotency model and can be cultured in different conditions: ground state (in serum-free medium with MEK and GSK3 inhibitors, or 2i embryonic stem cells [ESCs]), metastable state (with serum and leukemia inhibitory factor [LIF], or S/L ESCs), and primed state (in serum-free medium with bFGF and Activin A; or epiblast stem cells [EpiSCs]) [2, 3]. Each of these pluripotent states represents a downwards progression from the early preimplantation blastocyst to the post-implantation embryo. PSCs in 2i more closely resemble the pre-implantation ICM and produce chimeras upon blastocyst complementation more efficiently than metastable PSCs. On the contrary, EpiSCs resemble the post-implantation ICM and have negligible capacity to contribute to chimeras. The term metastable refers to the tendency of S/L ESCs to spontaneously oscillate between more naïve (ICM-like) conditions and the primed state [4]. Notably, cells in these three pluripotent states display differences in their epigenome, transcriptome and metabolome, as their function is controlled by different signalling pathways [5, 6]. Regarding metabolism, naive PSCs rely on both glycolysis and oxidative phosphorylation (through the tricarboxylic, TCA, cycle) for producing energy, whereas EpiSCs mostly rely on glycolysis [6, 7].

Besides providing energy, cell metabolism regulates myriad cellular functions comprehensively including modifications in DNA/histones, the transcriptome, and the proteome. Because metabolic features are variable in different cell types, changing metabolism can influence cell fate transitions. For instance, a rapid decrease of glycolysis during mouse ESC differentiation leads to reduced abundance of acetyl-coenzyme A (acetyl-CoA) and a consequent decline in histone lysine acetylation (an epigenetic mark associated with gene activation) at pluripotency loci [8]. Likewise, down regulation of SAM (S-adenosyl-methionine) caused by threonine depletion results in decreased trimethylation of histone H3 lysine 4, which slows PSC proliferation and facilitates differentiation [9]. Recently, a group of novel metabolites derived from short-chain fatty acids (*e.g*., propionyl-CoA, crotonyl-CoA, butyryl-CoA, and myristoyl-CoA, among others) have been shown to be substrate for lysine acylation of not only histones but also non-histone proteins involved in many cellular processes [10, 11]. These post-translational modifications are functionally relevant and distinct from protein lysine acetylation mediated by acetyl-CoA, further strengthening the link between metabolism and cellular functions beyond energy production. For example, during starvation histone 3 lysine 9 β-hydroxybutyrylation activates responsive genes in the mouse liver to induce adaption [12]. Similarly, lysine myristoylation of germline proteins can modulate the MPK-1/MAPK pathway to impact sex determination and reproductive development in *Caenorhabditis elegans* [13]. Moreover, lysine crotonylation (Kcr) in histones activates gene expression through yet unclear mechanisms, and a large repertoire of non-histone proteins can be crotonylated too [14, 15]. For example, crotonylation of RPA1 promotes homologous recombination-mediated DNA repair by enhancing its interaction with single-strand DNA [16]. Yet, comprehensive analysis of these novel post-translational modifications, including crotonylation, in different cell types and during cell fate transitions is largely lacking.

In this report, we have constructed an atlas of Kcr, with focus on non-histone proteins, in different mouse pluripotent states (2i ESCs, S/L ESCs, and EpiSCs) and also S/L ESCs allowed to exit pluripotency by removing LIF. We have identified 3628 high-confident Kcr sites on 1426 proteins among these four cell states. These sites reveal links between crotonylation and cell functions relevant for pluripotency maintenance, including RNA biogenesis, carbon metabolism, and the proteasome. Consistently, we observed that increased protein crotonylation caused by crotonic acid administration up-regulates pluripotency gene expression, delays differentiation, and enhances proteasome activity, a function relevant for pluripotency maintenance [17, 18]. Our findings provide a useful resource for understanding how metabolism regulates cell identity through crotonyl-CoA production, in particular the transition between different pluripotent states and differentiated cells.

## Results and discussion

### Quantitative lysine crotonylome analysis in different pluripotent states and differentiated cells

We aimed to investigate protein crotonylation in mouse PSCs in different states: 2i ESCs, S/L ESCs, EpiSCs, and also S/L ESCs triggered to exit pluripotency by LIF withdrawal for 4 days (hereafter referred to as differentiating cells) (**Figure 1A**). To monitor the different PSC states, we used ESCs and EpiSCs derived from crossed offspring of 129 female mice and OG2 transgenic male mice. These cells contain multiple copies of an *Oct4* distal enhancer-driven GFP reporter activated only in naïve pluripotency conditions [19, 20]. ESCs in 2i and S/L showed bright GFP fluorescence as well as the characteristic domed colony shape, whereas EpiSCs and differentiated cells had lost the GFP signal and displayed flat and flat/irregular colony shape, respectively (Figure S1A). To further verify the different cell identities, we performed reverse transcription-quantitative PCR (RT-qPCR) for general pluripotency markers (*Nanog* and *Oct4/Pou5f1*), pluripotent state-specific regulators (*Lin28a* and *Myc*) [21, 22] and EpiSC/differentiation markers (*Fgf5*). EpiSCs and differentiating cells showed higher expression of *Fgf5* compared to naïve ESCs (2i or S/L conditions), and 2i ESCs showed the lowest expression of *Myc, Lin28a*, and *Fgf5*, but the highest expression of *Nanog* and to a lesser extent *Oct4* (Figure S1B). These results demonstrate that our culture conditions truly represent the above-mentioned cell states.

**Figure 1.**
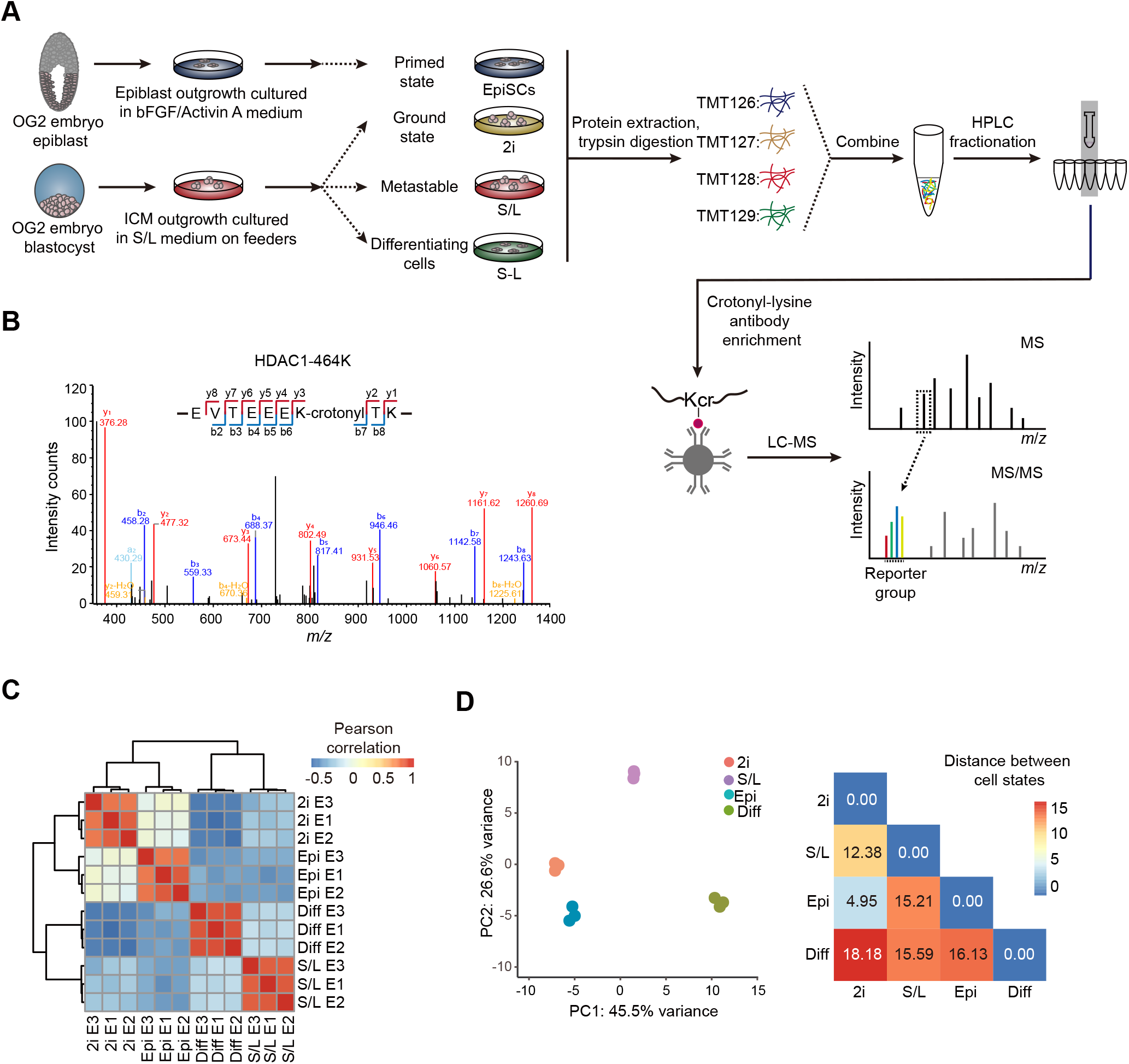
Generation of a high-confidence crotonylome map in three pluripotent states and differentiating cells. **A**. Schematic view showing the culture of cells in four different cell states and the workflow for the crotonylome profiling. **B**. Identification of HDAC1 crotonylation in LC-MS/MS; the MS/MS spectrum at m/z 1174.62 Da matches with the peptide –EVTEEEKTK-being crotonylated K7-crotonyl (68.023 Da). **C**. Hierarchical clustering of the normalized log2 intensity for crotonylated peptides of three LC-MS/MS experiments (biological replicates) in the four different cell states, colours in the heatmap indicate the pairwise Pearson correlation between the different samples (n = 3628). Epi: EpiSCs; Diff: Differentiating cells. **D**. Left: Principal component analysis of the crotonylome in the four different cell states; the variance was calculated using normalized log2 intensity. Right: Euclidean distances of crotonylation profiles between the indicated cell states.

Next, we quantified the lysine crotonylome of cells in these four culture conditions. To this end, we labelled the digested peptides from different conditions with tandem mass tags (TMT). We combined and fractionated the labelled peptides with high-performance liquid chromatography, and then enriched them with anti-Kcr antibodies [14] before conducting mass spectrometry analysis. Altogether, we identified 8,102 different sites in 2578 proteins among the four cell types, which were divided into high- (3628 sites in 1426 proteins) or low-confidence (4474 sites in 1152 proteins) candidates (Table 1 and S1A). High-confidence proteins were defined as quantified in at least two experiments out of three, and low-confidence as quantified only in one experiment). For example, we detected the chromatin regulator HDAC1 (Histone deacetylase 1), a known target of crotonylation [23], among the high-confidence crotonylated proteins (Figure 1B). In addition, we analyzed the total proteome (Table 1). Both the total proteome and the lysine crotonylome showed high reproducibility in three biological replicates (Figure S1C and 1C).

We then integrated our total proteome with the publicly available ‘core ESC-like gene module’ and the ‘adult tissue stem module’, which represent pluripotency regulators and differentiation regulators enriched in mouse ESCs and adult stem cells, respectively [24]. As anticipated, the ‘core ESC-like gene module’ scored the lowest in differentiating cells, whereas the ‘adult tissue stem module’ was the lowest in ground state PSCs but the highest in the differentiating cells (Figure S1D). Importantly, principal component analysis (PCA) of our crotonylome and proteome datasets could also distinguish cells in the four cell states (Figure 1D and S1G). The largest Euclidean distances in the crotonylome PCA were between 2i ESCs and the differentiating cells, consistent with the functional divergence between them. These observations support the notion that protein crotonylation may be functionally important for maintaining cell identity in these different culture conditions. In this regard, protein crotonylation is controlled by the balance between writers and erasers [25], whose changes in expression could explain putative differences in crotonylation between the four cell states. However, analysis of the total proteome showed that known crotonylation writers and erasers (also crotonylation readers) do not change noticeably between the four cell states (Figure S1E). Likewise, global crotonylation was at an equivalent level too (Figure S1F).

In summary, we have successfully profiled global protein crotonylation in different pluripotent states including 2i ESCs, S/L ESCs, EpiSCs, and also in early differentiating PSCs.

### Characteristics of the crotonylated proteins in the four cell states

We next studied the features of the 1426 high-confidence crotonylated proteins identified among the four cell states. Of note, 686 proteins showed at least two crotonylated sites, whereas 30 showed more than 12 modification sites (**Figure 2A** and Table S1B). Analysis of the subcellular localization using the COMPARTMENTS database [26] showed that these crotonylated proteins exert different functions in multiple compartments including nucleus, cytoplasm, plasma membrane, cytoskeleton, and mitochondria (Figure 2B and Table S1C). We also compared our complete Kcr dataset with published crotonylomes in different human cells including H1299 [15], HeLa [23], A549 [27], and HCT116 cells [28], and also peripheral blood from patients with kidney failure [29], all of which described identified crotonylated proteins without selection of high-confidence ones. Our dataset was the largest, with 886 out of the total 2578 crotonylated proteins (high- and low-confidence candidates) being unique to it (Figure 2C and Table S1D). To validate the identified candidates, we selected and overexpressed 15 high-confidence proteins and one low-confidence protein with a FLAG tag in HEK293T cells, followed by immunoprecipitation and western blotting with Kcr antibodies. The substrate for crotonyl-CoA generation, crotonic acid, was added to enhance the detection. Importantly, 11 out of 16 candidates showed increased basal Kcr signal, crotonic acid treatment boosted the basal signal in most of the selected candidates (15 out of 16) (Figure 2D and S2). Three out of the 15 validated proteins, including the pluripotency regulator LIN28A and the RNA m^6^A reader YTHDF2 [30], were exclusive to our own dataset. These results support the reliability of our proteomic experiments and analysis.

**Figure 2.**
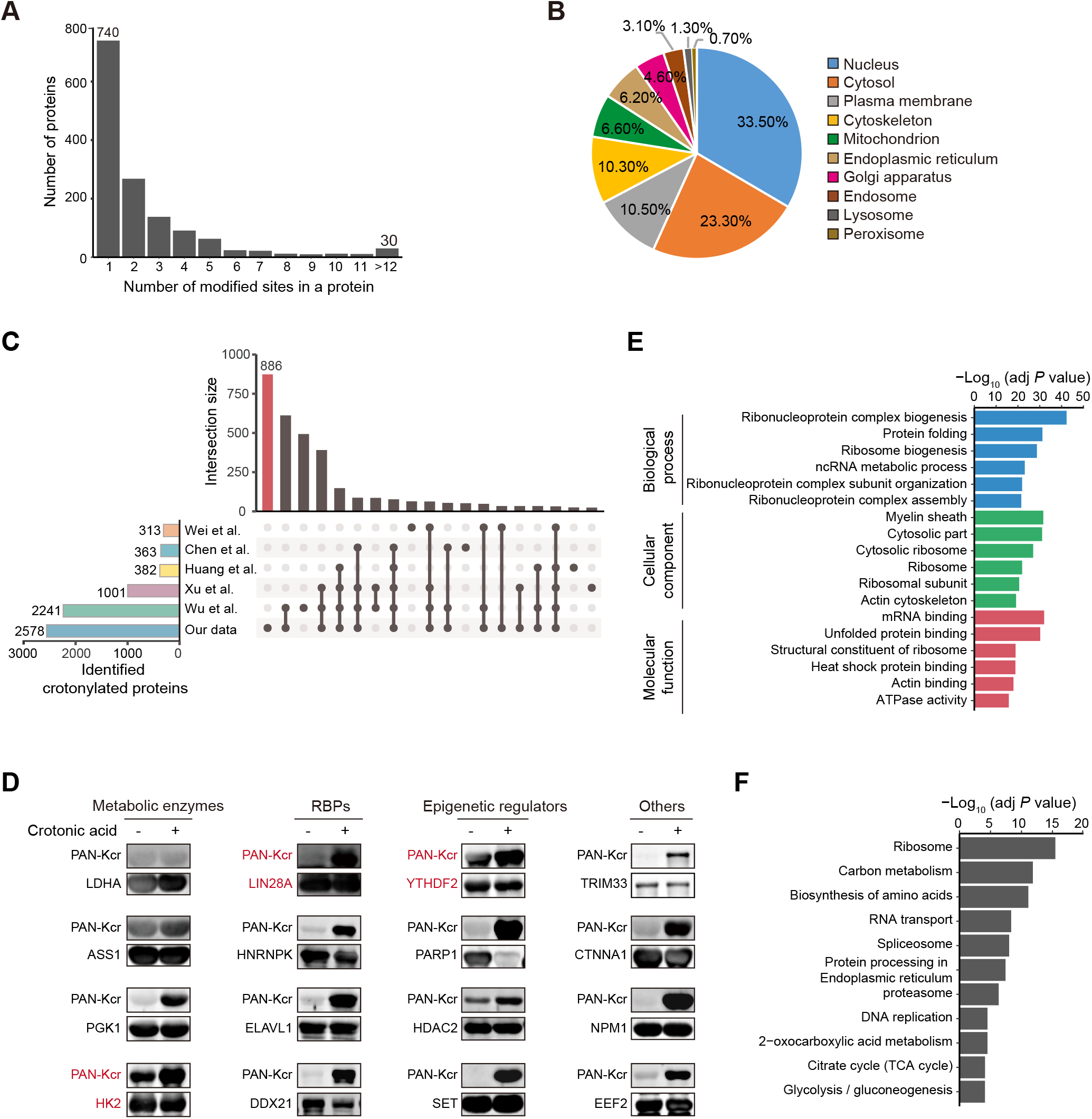
Characteristics of the crotonylated proteins identified in three pluripotent states and differentiating cells. **A**. Bar plot indicating the number of crotonylated sites in each high-confidence protein in our crotonylome datasets. **B**. Pie chart indicating the subcellular localization of the high-confidence crotonylated proteins. The localization was annotated using the COMPARTMENTS database in ‘knowledge’ channel. **C**. Comparison of our identified crotonylome data with crotonylome datasets from other published studies using human cells. **D**. Validation of protein crotonylation using ectopically expressed FLAG-tagged proteins in HEK293T cells with or without administration of 10 mM crotonic acid for 24 hours. Cell lysates were immunoprecipitated with anti-FLAG magnetic beads and analyzed by immunoblotting with anti-PAN Kcr antibodies. **E**. GO analysis of the high-confidence crotonylated proteins, the top 6 terms with smallest adjusted *P* value are shown (Fisher’s exact test, Benjamini-Hochberg corrected *P* < 0.01). **F**. KEGG analysis of the high-confidence crotonylated proteins, the top 11 terms with smallest adjusted *P* value are shown (Fisher’s exact test, Benjamini-Hochberg corrected *P* < 0.01).

### RNA binding proteins are highly represented in the four cell states crotonylome

We then performed gene ontology (GO) and KEGG pathway analysis of the 1426 high-confidence crotonylated proteins. Interestingly, GO analysis showed enrichment in RNA metabolism-related and ribosome-related terms (Figure 2E and Table S2A). Consistent with the former functional enrichment analysis, classical RNA binding domains (Figure S3A) and experimentally identified RNA binding proteins (RBPs) from other studies [31–33] (Figure S3B and Table S2B) were enriched in our high-confidence crotonylome dataset. This is perhaps not surprising because lysine is one of the most commonly enriched amino acids in the RNA binding domains of RBPs [34]. Thus, RBP crotonylation may be a widespread mechanism for regulating RNA-protein interactions by neutralizing the positive charge on lysine residues, in which case cell metabolism would appear as a major regulator of these interactions. This is particularly important for PSC function because many RBPs are essential regulators of pluripotency maintenance and exit [35]. For instance, K536 in the YTH domain of YTHDF2, which selectively recognizes the m^6^A modification on RNA [36], was identified as a high-confidence Kcr site. This may affect its binding affinity to m^6^A modified RNA and in turn regulate cell fate, as this RNA modification is necessary for pluripotency exit [37]. As for the ribosome, it is known that post-translational modifications of ribosome subunits influence protein translation [38], also suggesting a potential link of this function with cell metabolism. Notably, translational control has been implicated in pluripotency maintenance and differentiation [39, 40]. KEGG analysis showed enrichment in terms related to carbon metabolism and the proteasome (Figure 2F and Table S2A).

Because lysine residues are subjected to other posttranslational modifications in addition to crotonylation, we also compared our Kcr sites with reported acetylation, malonylation, succinylation, and ubiquitination mouse datasets from the Protein Lysine Modifications Database (PLMD) [41]. Our high-confidence Kcr sites significantly overlapped with other modifications, but a large number of Kcr sites were unique (Figure S3C and Table S2C). Moreover, we observed that the average amino acid distribution around crotonylated lysines is enriched in glutamic acid (E), in agreement with a previous crotonylome study [15] (Figure S3D). The negative charge of glutamic acid residues might synergize with the crotonylation-mediated suppression of lysine’s positive charge, changing the affinity of RBPs for substrates. Interestingly, it was reported that the glutamic acid-lysine (EK) rich region of the splicing factor SREK1/SRrp86 can function as an inhibitor of splicing [42]. Two lysine residues in this same domain of SREK1/SRrp86 were identified as low-confidence candidate sites in our dataset (Figure S3E and S3F).

Overall, these results demonstrate that protein crotonylation is a widespread lysine modification in mouse PSCs and early differentiating cells, affecting proteins with relevant functions in pluripotency maintenance and differentiation.

### Dynamics of protein crotonylation in the four cell states

We next compared the individual crotonylomes of the three pluripotent states and differentiating cells to gain insight into the potential functional consequences of differential protein crotonylation levels. For this purpose, we applied variance analysis (ANOVA with a cut-off of q < 0.01) and hierarchical clustering to the respective high-confidence crotonylated peptide datasets. We initially focused on non-histone proteins, which led to the identification of five categories (clusters I to V) among the four cell states (**Figure 3A** and B; Table S3A).

**Figure 3.**
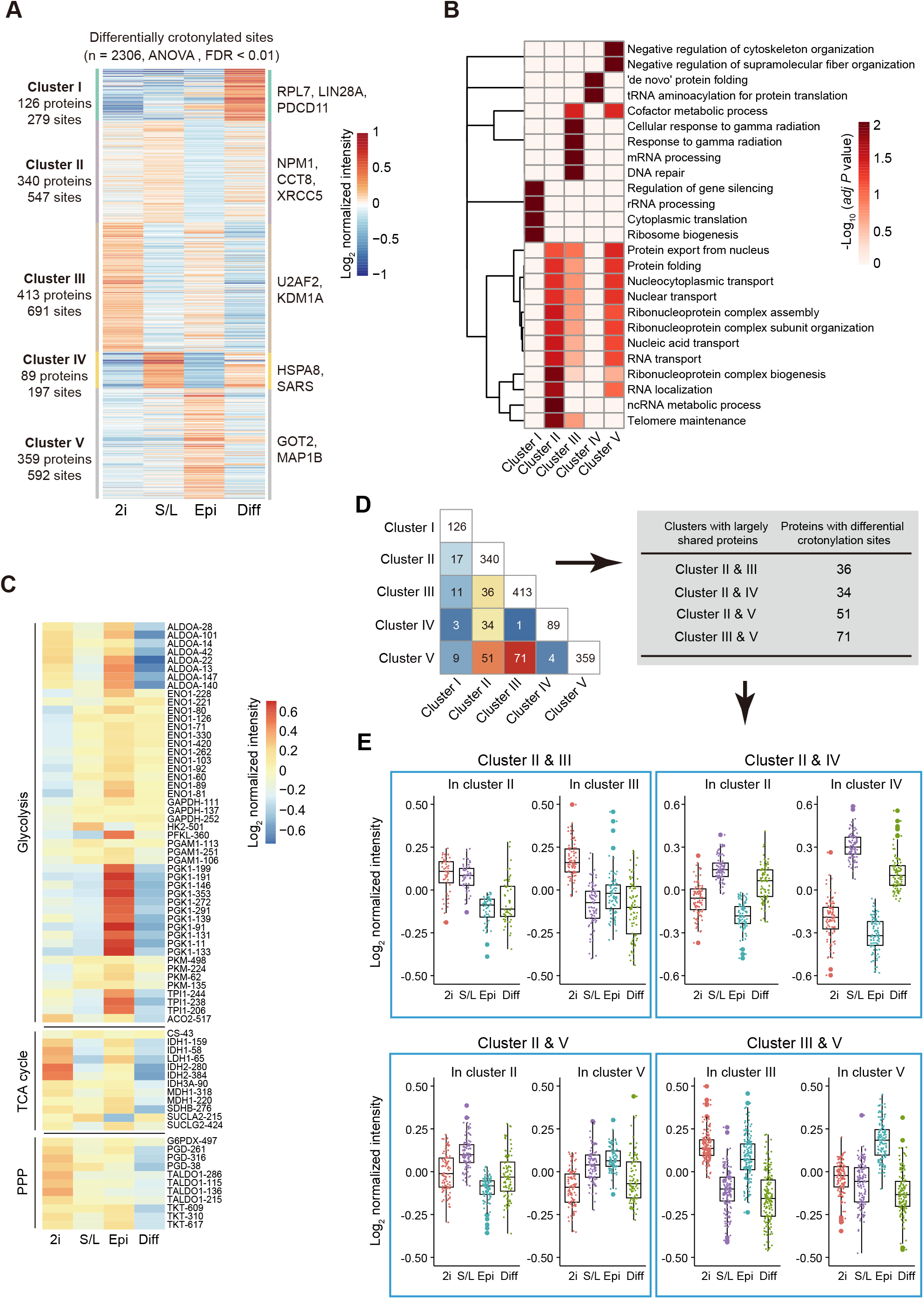
Functional clustering of the crotonylated sites in three pluripotent states and differentiating cells. **A**. Heatmap of crotonylated sites in high-confidence proteins shows distinct patterns in the four cell states (n = 2306, ANOVA test, FDR < 0.01). Normalized log_2_ intensity were grouped by hierarchical clustering into five clusters. **B**. GO analysis for all five clusters (Fisher’s exact test, Benjamini-Hochberg corrected *P* < 0.01, scaled without centering). **C**. Heatmap of differentially crotonylated enzymes involved in central carbohydrate metabolism (glycolysis, TCA cycle, and PPP pathway) (Fisher’s exact test, Benjamini-Hochberg corrected *P* < 0.01). **D**. Box plot indicating the number of crotonylated proteins overlapping between two clusters. Arrows indicate the direction of data processing. **E**. Crotonylation levels of specific lysine sites in the overlapped proteins between different clusters.

Cluster I corresponded to proteins highly crotonylated in differentiating cells compared to the three pluripotent states. This category included proteins related to protein translation including RPL7, LIN28A, and PDCD11. Cluster II included proteins displaying the lowest crotonylation level in EpiSCs, and was particularly enriched in proteins related to RNA localization/transport (*e.g*., NPM1), protein folding (*e.g*., CCT8), and telomere maintenance (*e.g*., XRCC5). Regarding the latter, crotonylation has been shown to facilitate telomere maintenance in chemically induced reprogramming [43], suggesting a potential cause-effect link. Cluster III corresponded to proteins that displayed the highest crotonylation level in 2i ESCs compared to the other three cell states, and included proteins related to mRNA processing (*e.g*., U2AF2) or DNA repair (*e.g*., KDM1A). Cluster IV consisted of proteins more highly crotonylated in S/L ESCs and differentiating cells compared to 2i ESCs and EpiSCs, and included regulators of protein folding (*e.g*., HSPA8) or tRNA aminoacylation (*e.g*., SARS). The last category, cluster V, were proteins more highly crotonylated in EpiSCs, and contained cofactors of metabolic processes (*e.g*., GOT2) and cytoskeletal regulators (*e.g*., MAP1B). We also observed a significant enrichment of ribonucleoprotein complex related terms in clusters III and IV, albeit to a lesser extent than in cluster II.

We looked in more detail into the enrichment of crotonylated peptides belonging to proteins involved in central carbon metabolism-related KEGG terms (TCA cycle, pentose phosphate pathway [PPP], and glycolysis) in the four cell stats. The crotonylation levels of these enzymes were highly dynamic among the different cell types (Figure 3C). Notably, we noticed enhanced crotonylation of glycolytic enzymes in EpiSCs compared to naïve ESCs, whereas TCA cycle and PPP enzymes were more enriched in 2i ESCs compared to EpiSCs. These data are consistent with increased glycolytic activity being a major route for energy production in EpiSCs and with the enhanced mitochondrial activity in naive ESCs compared to EpiSCs [6, 7]. This suggests that differential protein crotonylation in these cell states could actually be a contributing cause rather than a consequence of the observed metabolic differences. In this regard, it has been reported that acetylation of metabolic enzymes changes their function as an adaptive mechanism to environmental changes [44, 45].

We also observed that many crotonylated peptides were shared between clusters albeit with different intensity in each respective cluster (Figure 3D and E; Table S3B). For instance, the high-confidence K23 and K188 residues of the apoptosis regulator protein SET showed an opposed tendency between 2i ESCs and S/L ESCs, with K23 more crotonylated in 2i ESCs and K188 less (Figure S4A and S4B). This suggests: a) that crotonylation of different lysine residues of the same protein could have diverse functional consequences depending on the cell state, and b) that differential crotonylation of the same lysine in different cell states may be involved in causing distinctive functional features.

Because protein levels vary among the four cell states, we also performed the differential crotonylation analysis after calibrating the crotonylation level by protein abundance. Proteins with stable expression but that experience changes in their crotonylation status might be more critically involved in determining shifts in cell identity between the four cell states. After normalization, we identified 485 differential sites in 330 proteins between the four cell states (Figure S4C and S4D; Table S3C). These sites and proteins distributed in five clusters. Cluster I included proteins highly crotonylated in differentiating cells and was enriched in proteins related to the TCA cycle. Among these, we noticed for example ACLY, which participates in fatty acid synthesis to regulate pluripotency [46]. Cluster II consisted of proteins highly crotonylated in 2i PSCs and was enriched in GO terms related to mRNA processing, ribonucleoprotein complex assembly, and chromatin remodelling. The former GO terms included the classical RBP HNRNPU [47] and the deacetylase/decrotonylase HDAC1[48], which has been reported to regulate pluripotency. Cluster III represented proteins highly crotonylated in EpiSCs and contained proteins involved into RNA splicing and transport (*e.g*., DHX9). Cluster IV represented proteins displaying lower crotonylation in EpiSCs and it was only enriched for proteins related to protein folding (*e.g*., HSPA9). Cluster V contained proteins more crotonylated in EpiSCs compared to the other cell states. This cluster had higher crotonylation in EpiSCs than Cluster III; however, there were no enriched GO terms from these 20 proteins.

Histone crotonylation is highly dynamic during ESCs differentiation and is required for self-renewal [49]. Besides the analysis of non-histone proteins, we identified 25 crotonylation sites in histone proteins (Figure S5 and Table S3D). While extracting the histone for mass spectrometry might increase the number of identified histones crotonylation sites, our dataset is nevertheless valuable for studying the role of histone crotonylation in pluripotency and differentiation.

In summary, our data support that dynamic changes in protein crotonylation, which extend beyond the better studied histone targets, likely play relevant roles in controlling the transition between different pluripotent states and pluripotency exit.

### Crotonic acid enhances pluripotency gene expression associated with increased proteasome activity

Given the dynamic changes in protein crotonylation among the four cell states, we next asked whether modulating protein crotonylation would affect PSC pluripotency or differentiation. To test this, we treated S/L ESCs with crotonic acid. We observed significant upregulation of the pluripotency genes *Nanog* and *Dppa2*, and a concomitant downregulation of the differentiation genes *Fgf5* and *Pax6* compared to the control (**Figure 4A**). S/L ESCs treated with crotonic acid that were allowed to gradually differentiate upon LIF withdrawal for four days also exhibited higher pluripotency gene and lower differentiation gene levels than non-treated ESCs (Figure 4B), suggesting that enhanced protein crotonylation maintains pluripotency and slows pluripotency exit. It has been reported that crotonic acid treatment in mouse ESCs downregulates a group of pluripotency genes while upregulating two-cell (2C)-like state genes [43]. This state refers to cells closer to early post-fertilization time points, where embryonic cells have totipotent characteristics (can produce trophectoderm and extraembryonic mesoderm rather than only the cell types coming from the three embryonic germ layers). 2C-like cells are known to express lower levels of pluripotency factors than S/L or 2i PSC cultures from which they spontaneously originate [50]. The discrepancy with our data could be due to the use of different cell lines (N33 versus OG2 ESCs) or unnoticed variations in the culture conditions. Considering the very low number (~0.5%) of 2C-like cells in PSC cultures, single-cell gene expression studies would be necessary to fully clarify this. Regardless, the trend of the changes in both studies is in the same direction (increased potency).

**Figure 4.**
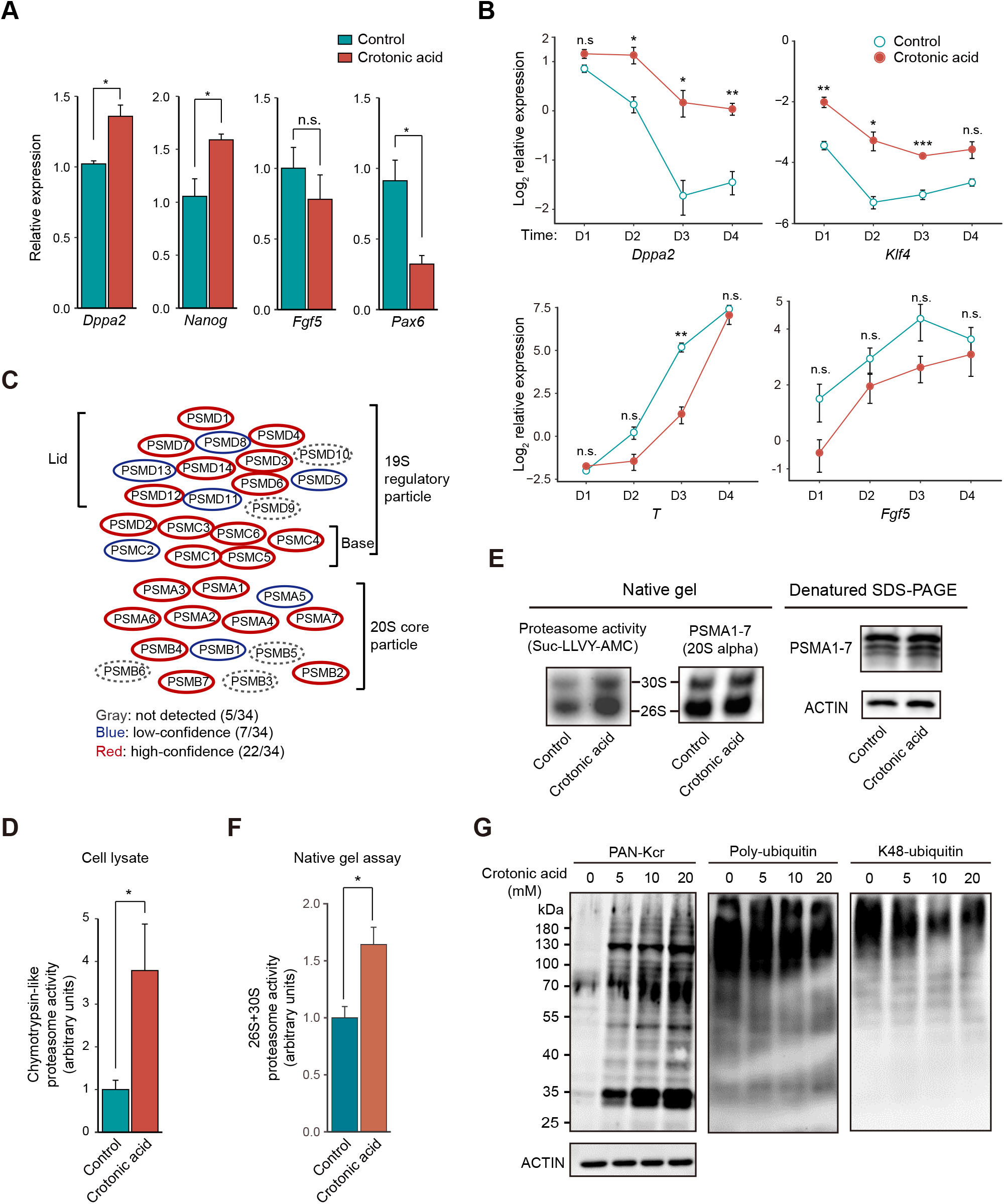
Crotonic acid promotes pluripotency and enhances proteasome activity. **A**. Relative expression of pluripotency and differentiation genes measured by RT-qPCR in ESCs cultured in S/L with or without 10 mM crotonic acid. Data are presented as the mean ± S.E.M. (n = 3 biological replicates with three technical replicates each, two-tailed unpaired Student’s *t*-test). \**P*□<□0.05, n.s.: not significant. **B**. Relative expression of representative pluripotency and differentiation genes measured by RT-qPCR in ESCs cultured in S/L and subjected to differentiation by withdrawing LIF over a time-course of 4 days. Data are presented as the mean ± S.E.M. (n = 3 biological replicates with three technical replicates each, two-tailed unpaired Student’s *t*-test). \**P*□<□0.05, \*\**P*□<□0.01, and \*\*\**P* < 0.001, n.s.: not significant. **C**, Schematic representation of the proteasome complex and the components identified in our crotonylome. Gray (dashed) ellipse: not detected; blue ellipse: low-confidence; red ellipse: high-confidence. **D**. Chymotrypsin-like proteasome activity measurement in ESCs cultured in S/L with or without 10 mM crotonic acid treatment for 48 hours (n = 4 biological replicates, two-tailed unpaired Student’s *t*-test). \**P*□ <□0.05. **E**. Chymotrypsin-like proteasome activity measurement in pre-purified proteasome by native gel electrophoresis, ESCs were cultured in S/L with or without 10 mM crotonic acid treatment for 48 hours. Samples were resolved by denatured SDS-PAGE and western blotting for analysis the 20S proteasome level and loading control. **F**. Quantified densitometry of the result shown in **E**, data are presented as the mean ± S.E.M. (n = 4 biological replicates, two-tailed unpaired Student’s *t*-test). \**P*<0.05. **G**. Representative western blotting of lysine crotonylation, poly-ubiquitination, and K-48 ubiquitination in ESCs cultured in S/L and untreated or treated with different concentrations of crotonic acid for 48 hours. ACTIN was the loading control.

Then, we sought to understand how crotonylation affects pluripotency. To this end, we first compared the crotonylated proteins in the four states with a reported ESC-specific gene set (genes highly expressed in mouse/human ESCs) [33]. This showed that 466 ESC-specific gene-encoded proteins are crotonylated (Figure S6A). GO cellular component analysis of these proteins showed enrichment in terms related to ‘proteasome complex’ as well as ‘peptidase complex’, among others (Figure S6B and Table S4A). Indeed, most of the proteasome complex subunits were crotonylated (29/34 total and 22/34 high-confidence) in our dataset (Figure 4C and Table S4B). It has been shown that proteasome activity is positively associated with pluripotency maintenance [17, 18], and our crotonylated dataset included the two proteasomal subunits, PSMD11 and PSMD14, reported as pluripotency regulators. Thus, we tested chymotrypsin-like proteasome activity in S/L ESCs treated or untreated with crotonic acid. Adding crotonic acid significantly enhanced proteasome activity compared to the control in crude lysate (Figure 4D) and also in the native gel assay (Figure 4E and 4F). The latter experiment can separate active proteasome complexes from other biomacromolecules based on the molecular weight [51]. Moreover, western blotting showed that the increase of protein crotonylation signals caused by adding crotonic acid associate with a decrease of poly-ubiquitin and K48-ubiquitin signals in cell lysates (Figure 4G). This further points to an increase in proteasome activity after crotonic acid treatment.

Taken together, these experiments suggested a link between crotonylation of proteasome subunits in PSCs and pluripotency maintenance. Additional work will be necessary to clarify whether this link is causal and how it relates to the crotonylation of other proteins involved in pluripotency control including histones [14].

## Conclusions

Our global atlas of protein crotonylation in four different pluripotent cell states has identified targets essential for pluripotency regulation. Among these, it is worth noting factors involved in RNA biogenesis, protein translation, metabolic factors, and proteasome subunits. Further work will be important to validate the specific impact of crotonylation on the function of these proteins, and also how these effects crosstalk with the consequences on gene expression of histone crotonylation [49]. Hence, our dataset will be an important resource for future studies on pluripotency and early pluripotency exit. Our results may also be helpful to understand how changes in metabolism influence cell function in other contexts through variation in the levels of crotonyl-CoA.

## Materials and methods

### Isolation of ESCs and EpiSCs

OG2 ESCs were isolated in previous study [52]. For derivation of OG2 EpiSCs, the late epiblast layer was dissected intact from a pre-gastrulation stage mouse blastocyst [E5.75], maintained in FA medium (DMEM/F12 [Catalog No. SH30023.01B, Hyclone, Buckinghamshire, United Kingdom] and Neurobasal medium [Catalog No. 21103049, Gibco, Carlsbad, CA] mixed 1:1, supplemented with 1× non-essential amino acids [Catalog No. 11140050, Gibco], 1× GlutaMAX [Catalog No. 35050061, Gibco], 1 mM sodium pyruvate [Catalog No. 11360070, Gibco], 50 U/ml penicillin/streptomycin [Catalog No. SV30010, Hyclone], 1× N-2 [Catalog No. 17502048, Gibco], 1× B-27 [Catalog No. 17504044, Gibco], 12.5 ng/ml bFGF [Catalog No. 233-FB-050, R&D, Minneapolis, MN], and 20 ng/ml Activin A [Catalog No. 338-AC-050, R&D]) on Matrigel (Catalog No. 354248, Corning, Corning, NY)-coated plates and passaged 4-8 days after isolation.

### ESC and HEK293T culture

OG2 ESCs were cultured in the presence of 1× non-essential amino acids, 1× GlutaMAX, 1 mM sodium pyruvate, 50 U/ml penicillin/streptomycin, 1000 U/ml LIF (Catalog No. ESG1107, Millipore, Darmstadt, Germany), in either DMEM-high glucose medium (Catalog No. SH30022.01, Hyclone) supplemented with 15% fetal bovine serum (Catalog No. S1580 Biowest, Nuaillé, France) on feeder (mitomycin-treated mouse embryonic fibroblasts)-coated plates (for the S/L condition) or DMEM/F12 and Neurobasal medium mixed 1:1, supplemented with 1× N-2, 1× B-27, 1 μM PD0325901, and 3 μM CHIR99021 on 0.1% gelatin (Catalog No. ES-006-B, Millipore)-coated plates (for the 2i condition). For differentiating cells, ESCs were cultured in the same medium as S/L ESCs but without LIF and on 0.1% gelatin-coated plates for four days. HEK293T cells were cultured in DMEM-high glucose supplemented with 10% fetal bovine serum.

### Protein extraction and trypsin digestion for LC-MS/MS

Cells were lysed on ice in a buffer containing 8 M urea, 10 mM DTT, 50 mM nicotinamide, 3 μM TSA, and 1× protease inhibitor cocktail (Catalog No. 11697498001, Roche, Basel, Switzerland). Lysates were centrifuged (20,000 g) at 4 °C for 10 minutes. Supernatants were precipitated with cold 15% trichloroacetic acid at −20 °C for 2 hours, following 20,000 g centrifugation at 4 °C for 10 minutes. The precipitated proteins were dissolved in a pH 8.0 buffer containing 8 M urea and 100 mM triethylammonium bicarbonate buffer. The protein solution was reduced with 10 mM DTT at 37 °C for 1 hour followed by alkylation with 20 mM iodoacetamide at room temperature for 45 minutes protected from light. The alkylated protein samples were diluted by adding 100 mM triethylammonium bicarbonate buffer. Trypsin (Catalog No. V5111, Promega, Madison, WI) was added at 1:50 (w/w) trypsin to protein ratio overnight, and then at 1:100 ratio for 4 hours.

### TMT labelling

The digested proteins were desalted by running on a Strata X C18 SPE column (Catalog No. 8B-S100-AAK, Phenomenex, Torrance, CA) and then vacuum-dried. Desalted peptides were labelled with TMTsixplex™ Isobaric Label Reagent Set (Catalog No. 90061, ThermoFisher Scientific, Waltham, MA) following the manufacturer’s protocol. Briefly, TMT was added at 1:1 (U/mg) TMT reagent to protein ratio for 2 hours at room temperature and then the samples were desalted.

### Peptide fractionation

TMT-labelled peptides were fractionated by high pH reverse-phase HPLC using an ZORBAX Extend-C18 column (Catalog No. 770450-902, Santa Clara, CA) (5 μm particles, 4.6 mm inner diameter, 250 mm length). Briefly, labelled peptides were separated with a gradient of 2-60% acetonitrile in 10 mM ammonium bicarbonate into 80 fractions for 80 minutes. Fractionated peptides were combined into 18 fractions for total proteome analysis or 8 fractions for crotonylome analysis.

### Kcr enrichment

Kcr-containing peptides were dissolved in pH 8.0 NETN buffer (100 mM NaCl, 1 mM EDTA, 50 mM Tris-HCl, and 0.5% NP-40). Dissolved peptides were incubated with antiKcr antibody-coated agarose beads (Catalog No. PTM-503, PTM Biolabs, Hangzhou, China) overnight at 4 °C. Beads were then washed with NETN buffer and bound peptides were eluted with 0.1% trifluoroacetic acid. The resulting peptides were desalted with C18 ZipTips (Catalog No. ZTC18S008, Millipore) before LC-MS/MS analysis.

### LC-MS/MS analysis

Enriched peptides were dissolved in 0.1% formic acid and loaded onto a home-made reverse-phase analytical column (15 cm length, 75 μm inner diameter). Peptide samples were eluted with a gradient elution system: an increasing gradient of solvent A (0.1% formic acid in 98% acetonitrile) from 7% to 20% over 24 minutes, 20% to 35% in 8 minutes and climbing to 80% in 5 minutes then holding at 80% for the last 3 minutes, all at a constant flow rate of 300 nl/minute on an EASY-nLC 1000 UPLC system (Catalog No. LC120, ThermoFisher Scientific). The eluted peptide samples were analyzed by a Q Exactive Plus™ hybrid quadrupole-Orbitrap™ mass spectrometer (Catalog No. IQLAAEGAAPFALGMAZR, ThermoFisher Scientific). The electrospray voltage was set to 2.0 kV. The m/z scan range was fixed from 350 to 1800 for full scan, and intact peptides were detected at a resolution of 70,000. Peptides were then selected for MS/MS using NCE setting as 30 and the fragments were detected at a resolution of 17,500. Data-dependent acquisition was used for MS data collection.

### LC-MS/MS data analysis

The resulting TMT data were processed using MaxQuant (v.1.4.1.2) with integrated Andromeda search engine [53]. Tandem mass spectra data were searched against non-redundant mouse protein amino acid sequence from Uniprot databases (https://www.uniprot.org/) concatenated with reverse decoy database. Trypsin/P was selected as cleavage enzyme permitting up to two missing cleavages per peptide for total proteome analysis or four missing cleavages per peptide for crotonylome analysis. Mass error was set to 10 ppm for precursor ions and 0.02 Da for fragmented ions. Carbamidomethylation on cysteine was specified as fixed modification, oxidation on methionine, crotonylation on lysine, and acetylation on the N-terminal of the protein were specified as variable modifications. FDR threshold for proteins, peptides, and modification sites were set at less than 1%. Minimum peptide length was set at 7. For peptide quantification, TMT6plex was selected. All other parameters in MaxQuant were used default setting.

### Statistical analysis

All statistical analyses were performed using R (3.5.1). For protein homology analysis, protein sequences in identified human crotonylated proteins datasets [15, 23, 27–29] were inputted into ‘blastp’ function in blast+ (2.8.1) with the mouse crotonylated proteins we discovered. Resulting homologous proteins with *E* value < 1*e*-10 were considered as overlapping proteins in the human crotonylated proteins datasets. The ‘upset’ function in R package “UpSetR” (1.4.0) was used to visualize the number of identified crotonylated proteins and intersections of this study with the various human crotonylated protein datasets. Pairwise experimental Pearson correlation matrix was generated using ‘cor’ function in R with the option “use = all.obs”. Correlation matrix was clustered using hierarchical clustering by Euclidean distance and complete linkage method. Cell state variation was defined by performing PCA with ‘prcomp’ function in R. For analysis of the differentially modified sites, we used ANOVA test to generate *P* values for each high-confidence peptide. The generated *P* values were corrected by multiple hypotheses with FDR (Benjamini-Hochberg). Pearson correlation results and expression data were visualized using the ‘pheatmap’ function within the R package “pheatmap” (1.0.12).

### Cellular compartment analysis

Subcellular localization of proteins was analyzed using the COMPARTMENTS database [26] in ‘knowledge’ channel. Only the major compartments in eukaryotic cells (nucleus, cytosol, plasma membrane, cytoskeleton, mitochondrion, endoplasmic reticulum, Golgi apparatus, endosome, lysosome, and peroxisome), with possibility score > 1, were selected.

### GO and KEGG annotation

GO and KEGG annotations were performed using the R package “clusterProfiler” (3.10.1) [54] with the ‘enrichGO’ and ‘enrichKEGG’ function, respectively.

### Co-modification analysis

For analysis of crosstalk between crotonylation sites and other lysine modifications, we used the PLMD database [41], which lists 20 types of lysine modifications in multiple species. Co-modified sites were generated by merging our dataset with mouse PLMD dataset.

### Protein domain analysis

Protein domains of crotonylated proteins were annotated using InterProScan (http://www.ebi.ac.uk/interpro/) based on protein sequence alignment method with default settings. Fisher’s exact test and standard FDR control method were used to test the enrichment of the annotated domains from crotonylated proteins against proteome-wide domains.

### Crotonylation motif analysis

The frequency of amino acids surrounding crotonylated lysine residues was generated using iceLogo [55]. Briefly, high-confidence Kcr-central peptide sequences (6 amino acids upstream and downstream of the modified site) were used as input against the precompiled *mus musculus* protein sequences from Swiss-Prot, the percent difference was set as scoring system with a significance cut-off of *P* = 0.05.

### Validation of crotonylated proteins by co-immunoprecipitation

HEK293T cells were transfected with plasmids producing FLAG-tagged proteins. After transfection, cells were lysed with TNE lysis buffer (50 mM Tris-HCl pH 7.5, 150 mM NaCl, 0.5% NP-40, 1 mM EDTA, 10 mM sodium butyrate, and 1× protease inhibitor cocktail). Lysates were sonicated with a Bioruptor (Catalog No. B01020001 Diagenode, Ougrée, Belgium) sonicator (low power, 30 second ON/OFF, 5 cycles), followed by 10,000 g centrifugation at 4 °C for 10 minutes to remove the undissolved particles. Then, they were incubated with pre-washed anti-FLAG M2 magnetic beads (Catalog No. M8823, Sigma, Darmstadt, Germany) at 4 °C overnight with rotation, and eluted by heating at 95 °C for 5 minutes with SDS-PAGE loading buffer. The following primary antibodies were used for immunoblotting: anti-FLAG (Catalog No. F7425, Sigma) and anti-Kcr (Catalog No. PTM-501, PTM Biolabs).

### Western blotting

Western blotting was performed using standard procedure after lysing cells with a buffer containing 10 mM Tris-HCl pH 7.4, 10 mM EDTA, 50 mM NaCl, 1% Triton X-100, 0.1% SDS, 10 mM N-ethylmaleimide, and 1× protease inhibitor cocktail. Lysates were boiled for 10 minutes to ensure inactivation of deubquitinase enzymes. Denatured lysates were sonicated with a Bioruptor sonicator (low power, 30 second ON/OFF, 5 cycles), followed by 10,000 g centrifugation at 4 °C for 10 minutes to remove undissolved particles. Lysates were then subjected to immunoblotting with the following primary antibodies: anti-polyubiquitin (Catalog No. 14220, Cayman, Ann Arbor, MI), anti-k48-linkage specific polyubiquitin (Catalog No. 8081S, CST, Danvers, MA), anti-NANOG (Catalog No. A300-397A, Bethyl, Montgomery, MA), and anti-β-ACTIN (Catalog No. A2228, Sigma).

### Proteasome activity assay

Chymotrypsin-like proteasome activity was measured using a Proteasome Activity Assay Kit (Catalog No. ab107921, Abcam, Cambridge, United Kingdom) following the manufacturer’s protocol. In brief, cells with or without crotonic acid treatment were collected in 0.5% NP-40 and homogenized by pipetting up and down a few times. After 10,000 g centrifugation at 4 °C for 10 minutes, the supernatants were incubated with proteasome substrate (Suc-LLVY-AMC) at 37 °C for 20 minutes. The released free AMC (7-amino-3-methylcoumarin) fluorescence was detected on a microplate fluorometer (350 nm excitation, 440 nm emission) after 30 minutes and then 60 minutes at 37 °C. Background signal was corrected by subtracting the 30-minute reading from the 60-minute reading and data were then normalized against the control sample.

### Native gel assay for proteasome activity

Native gel assay for proteasome activity was performed according to the previous publication [17]. Briefly, the cells with or without crotonic acid treatment were collected in proteasome activity assay buffer (50 mM Tris-HCl pH 7.5, 5 mM MgCl2, 5 mM ATP, 1 mM DTT, and 10% glycerol) and lysate by passing 10 times through the syringe with 27-G needled. After 12,000 g centrifugation at 4 °C for 10 minutes, the supernatants were loaded on 3–12% NativePAGE Bis-Tris gel (Catalog No. BN1003BOX, Invitrogen, Carlsbad, CA) in NativePAGE running buffer (Catalog No. BN2001, Invitrogen) containing 5 mM MgCl_2_ and 1 mM ATP, and run at 4 °C for 150 min at 150 V. The native gel was then incubated with 300 μM Suc-LLVY-AMC (Catalog No. HY-P1002, MedChemExpress, Monmouth Junction, NJ) diluted in NativePAGE running buffer. Chymotrypsin-like proteasome activity was detected on a FluorChem E system (Catalog No. 92-14860-00, ProteinSimple, San Jose, CA) in UV channel with 460 nm filter. The gel was then incubated with transfer buffer with 1% SDS for 10 minutes followed by incubation with transfer buffer for 10 minutes. The denatured gel was then transferred to PVDF membrane and incubated with anti-Proteasome 20S alpha antibody (Catalog No. ab22674, Abcam).

## Ethical statement

Animal experiments in this study were compliant with all relevant ethical regulations for animal research, and conducted under the approval of the Animal Care and Use committee of the Guangzhou Institutes of Biomedicine and Health under licence number 2007007.

## Data availability

Mass spectrometry proteomic data have been deposited to the ProteomeXchange Consortium via the PRIDE [56] partner repository with the dataset identifier PXD017121. MS/MS spectrum data of all identified crotonylated peptides have been deposited to the ProteinProspector database. They can be accessed in http://msviewer.ucsf.edu/prospector/cgi-bin/msform.cgi?form=msviewer by the following search keys: Replicate 1: zxlfv5pcbb; Replicate 2: g7bxkypz5c; Replicate 3: rdqvecojfj.

## Authors’ contributions

ZC and XB conceived the original idea and YL contributed to it. YL performed most of the experiments, CB performed the LC-MS/MS experiments, JM, JH, and MJ contributed to the experiments, YL performed the bioinformatic analyses with help from CW. MAE contributed to all data analysis. LG isolated the EpiSC cell line under the supervision of JC. XB, ZC, and MAE provided financial support and supervised the project. MAE, XB, and YL wrote the manuscript. CB, CW, and GV helped to revise the manuscript. All authors read and approved the final version of the manuscript.

## Acknowledgements

We thank all members of the Laboratory of Integrative Biology for their support. We also thank Xiaofei Zhang, from the Guangzhou Institutes of Biomedicine and Health, for constructive advice on proteomic data analyses. This work was supported by the National Key Research and Development Program of China (2016YFA0100102, 2016YFA0100701, and 2018YFA0106903), the Strategic Priority Research Program of the Chinese Academy of Sciences (XDA16030502), the Natural Science Foundation of Guangdong Province (2018B030306042), the Youth Innovation Promotion Association of the Chinese Academy of Sciences (2015294), and an Innovation Team Project grant from the Guangzhou Regenerative Medicine and Health Guangdong Laboratory (2018GZR110103001). Carl Ward is supported by a Zhujiang Talent-Overseas Postdoctoral Funding Grant and a President’s International Fellowship Initiative grant from the Chinese Academy of Sciences.

---

**Table 1 Summary of proteome and crotonylome profiling results.**

## Supplementary material

**Supplementary Figure S1.**
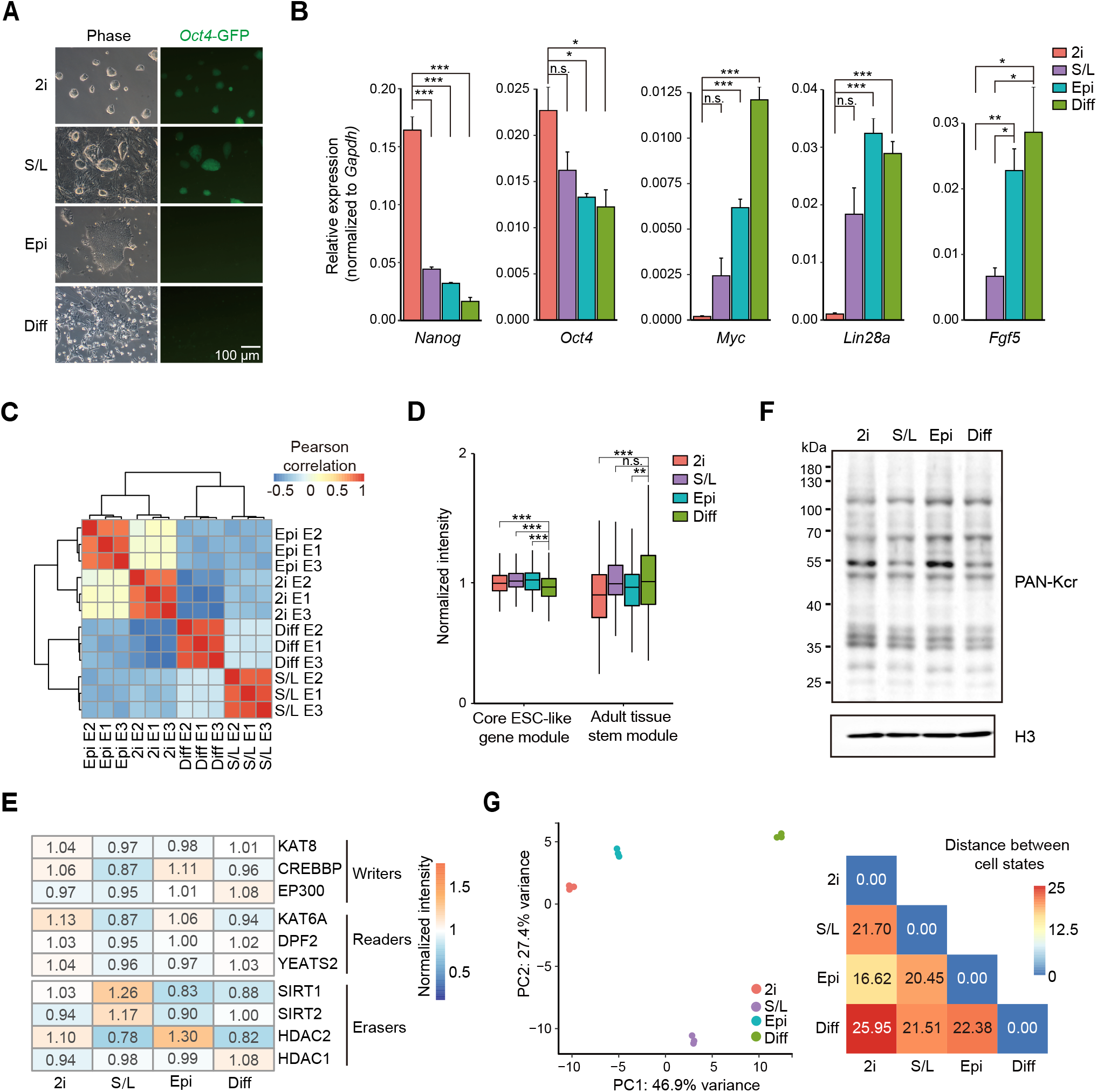
Characterization of different cell states. *Related to Figure 1*. **A**. Representative images showing PSCs cultured in different conditions, bright-field (phase) and green fluorescence for the same fields are shown. Scale bar = 100 μm. **B**. Relative expression of selected pluripotency and differentiation genes measured by RT-qPCR in the four different cell states. Data are presented as mean ± S.E.M. (n = 3 biological replicates with 3 technical replicates each, two-tailed unpaired Student’s *t*-test). \**P*□<□0.05, \*\**P*□<□0.01, and \*\*\**P* < 0.001, n.s.: not significant. *Gapdh* was used as the housekeeping control gene. **C**. Hierarchical clustering of the normalized log_2_ intensity for the total proteome of three different LC-MS/MS experiments (biological replicates) in four different cell states, colours in the heatmap indicate pairwise Pearson correlation between the different datasets (n = 4728). **D**. Normalized log2 intensity of ‘Core ESC-like gene module’ and ‘Adult tissue stem module’ gene sets from MSigDB in the four different cell states (n = 3 biological replicates, two-tailed unpaired Student’s *t*-test). \*\**P*□<□0.01, and \*\*\**P* < 0.001, n.s.: not significant. **E**. Heatmap showing the protein expression levels of crotonylation regulators in our total proteome analysis. **F**. Western blotting showing the global protein crotonylation levels in the four different cell conditions. **G**. Left: PCA of the total proteome in the four different cell states is shown; the variance was calculated using normalized log2 intensity. Right: Euclidean distances of proteome profiles between the indicated cell conditions.

**Supplementary Figure S2.**
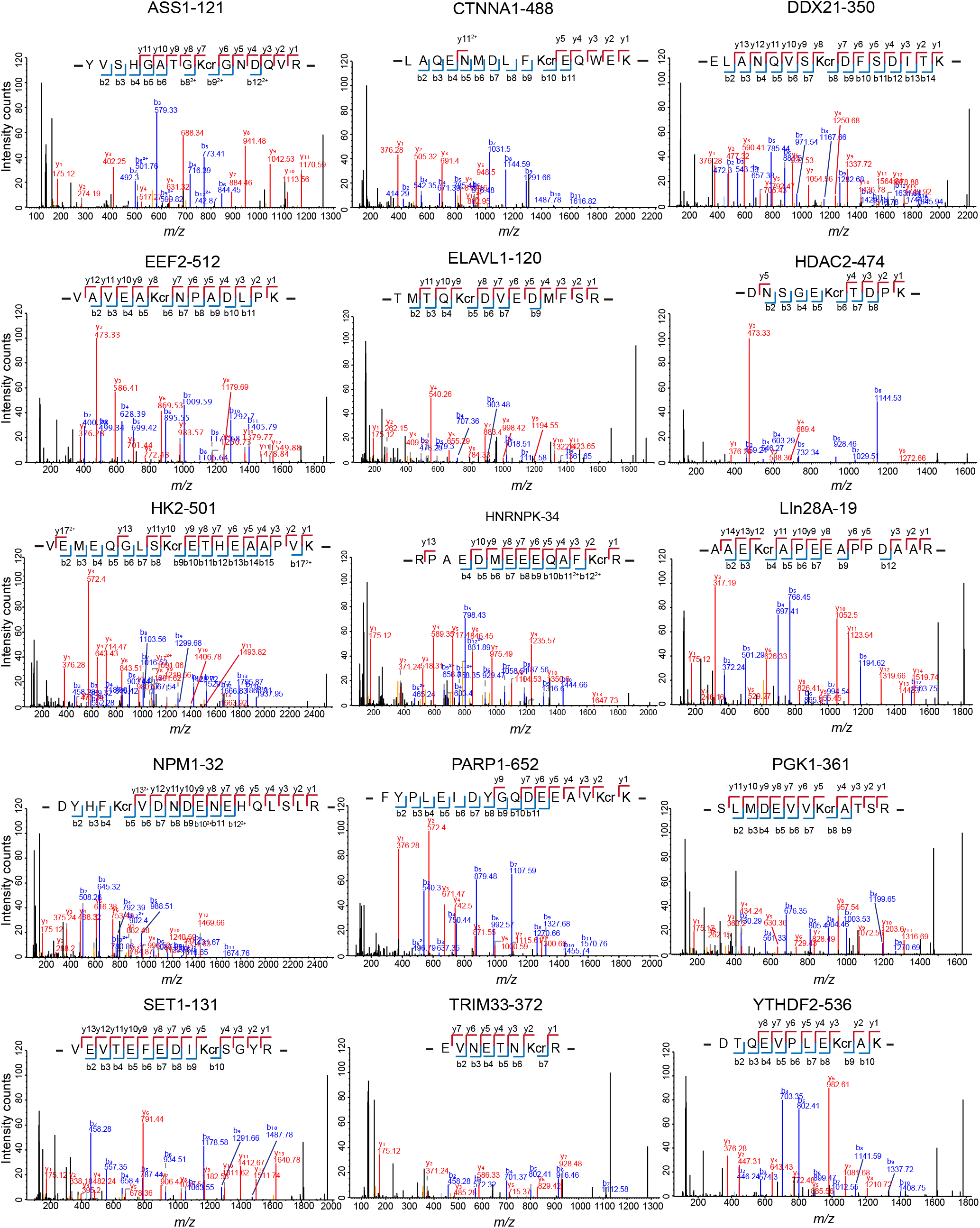
MS/MS spectrum of identified Kcr peptides of the 15 western blotting-validated proteins in Figure 2D.

**Supplementary Figure S3.**
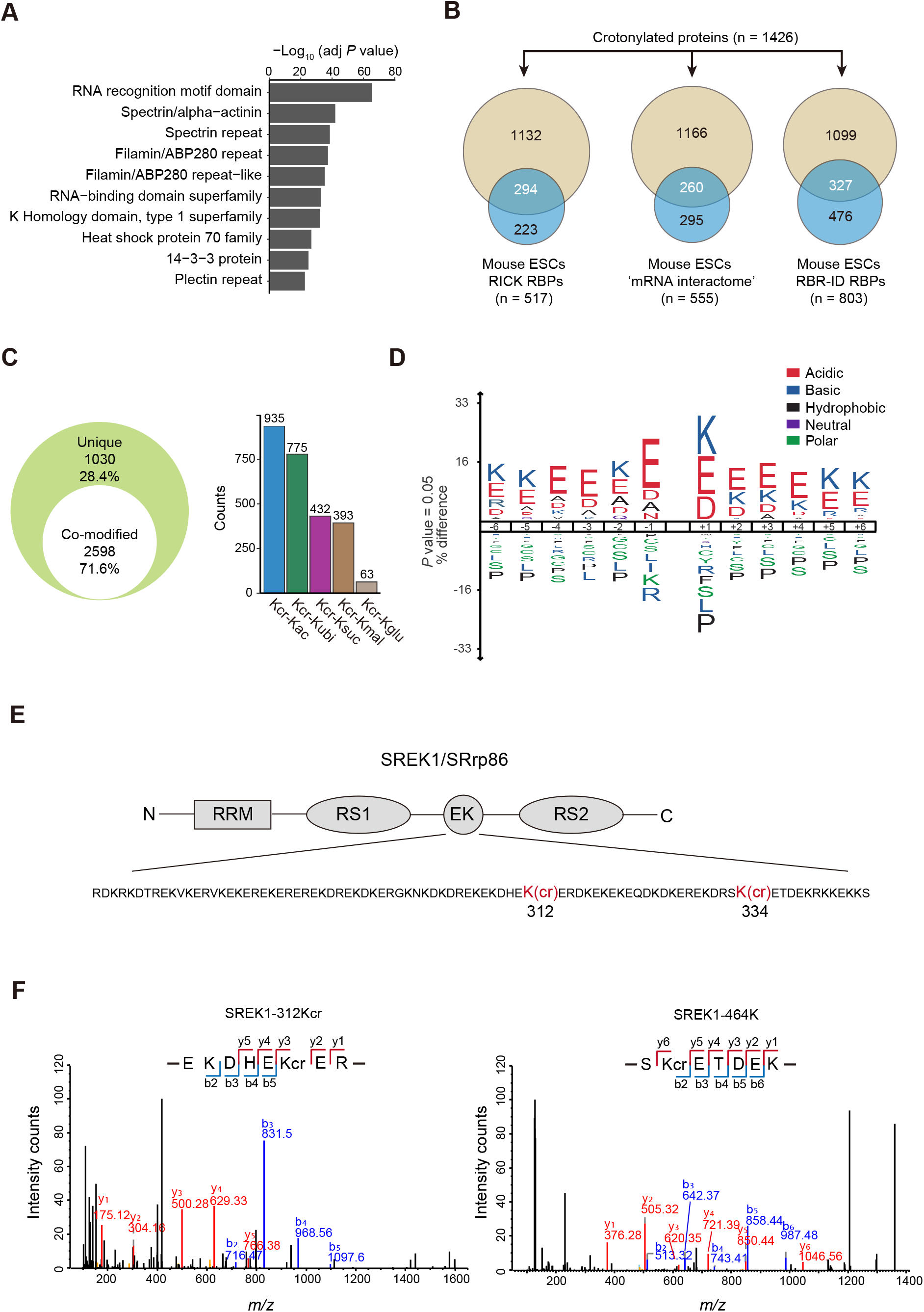
Functional characterization of the crotonylome in different cell states. *Related to Figure 2*. **A**. Protein domain enrichment analysis of the high-confidence crotonylated proteins, the top 10 terms with the smallest adjusted *P* value are shown (Fisher’s exact test, Benjamini-Hochberg corrected *P* < 0.01). **B**. Venn diagrams comparing our crotonylated datasets with published mouse ESCs RBP datasets. **C**. Comparison of the numbers of the overlapped lysine sites between our high-confidence crotonylome and other lysine modification datasets in the PLMD database. Ac: acetylation; ubi: ubiquitination; suc: succinylation; mal: malonylation; glu: glutarylation. **D**. Consensus sequence motif extracted from all high-confidence Kcr sites in our datasets. **E**. Schematic showing lysine crotonylation sites in the EK rich region of SREK1/SRrp86. **F**. MS/MS spectrum of K321cr (left) and K334cr (right) for SREK1/SRrp86.

**Supplementary Figure S4.**
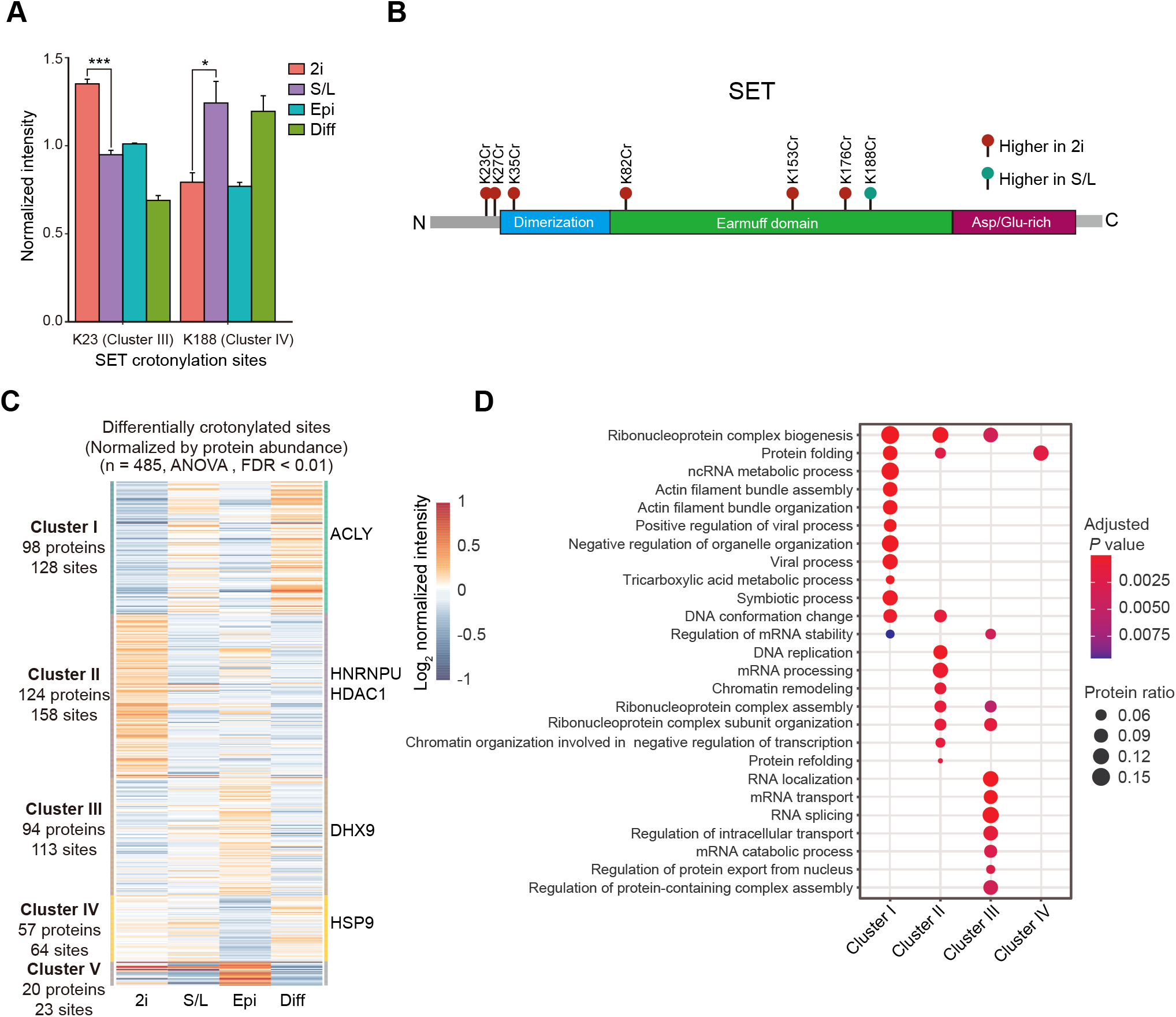
Crotonylation of SET protein in different cell states. *Related to Figure 3*. **A**. Crotonylation levels of two lysine sites of SET protein in the four different cell states. Data are presented as the mean ± S.E.M. (n = 3 biological replicates, two-tailed unpaired Student’s *t*-test). \**P*□ <□0.05, and \*\*\**P* < 0.001. **B**. Schematic showing the high-confidence Kcr sites of SET. Green circle: higher crotonylation level in S/L ESCs than in 2i ESCs; red circle: higher crotonylation level in EpiSCs than in 2i ESCs. **C**. Heatmap of differential crotonylated sites in the four different cell states after being normalized by protein abundance (n = 485, ANOVA test, FDR < 0.01). **D**. GO biological process analysis for the different clusters of proteins in **C** (Fisher’s exact test, Benjamini-Hochberg corrected P < 0.01).

**Supplementary Figure S5.**
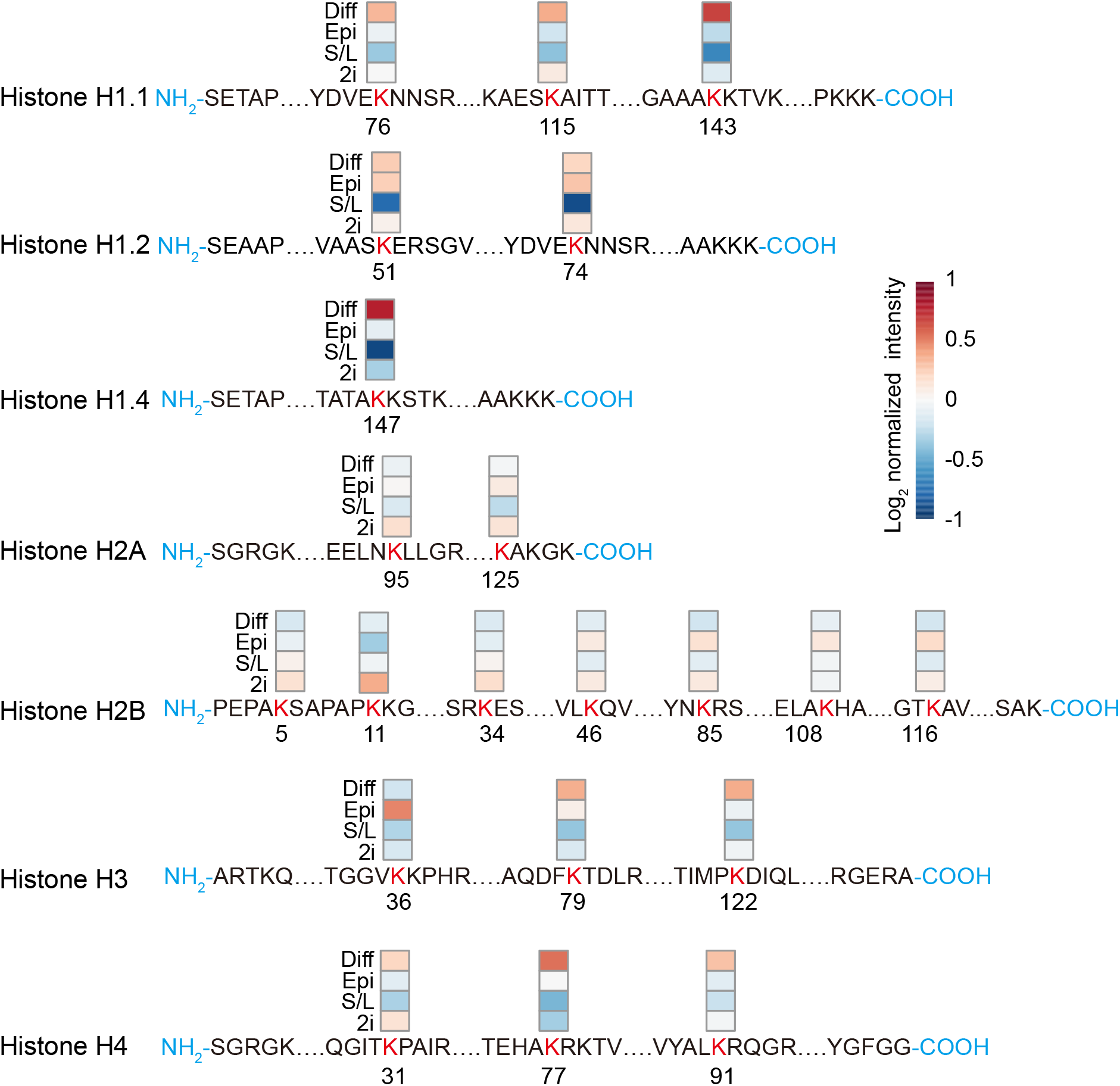
Histone crotonylation analysis in different cell states. *Related to Figure 4*. Diagrams showing the quantitative information of histone Kcr sites in the four cell states of our study. For the isoform-specific histone sites that occurred at the same position, the mean normalized intensity has been used.

**Supplementary Figure S6.**
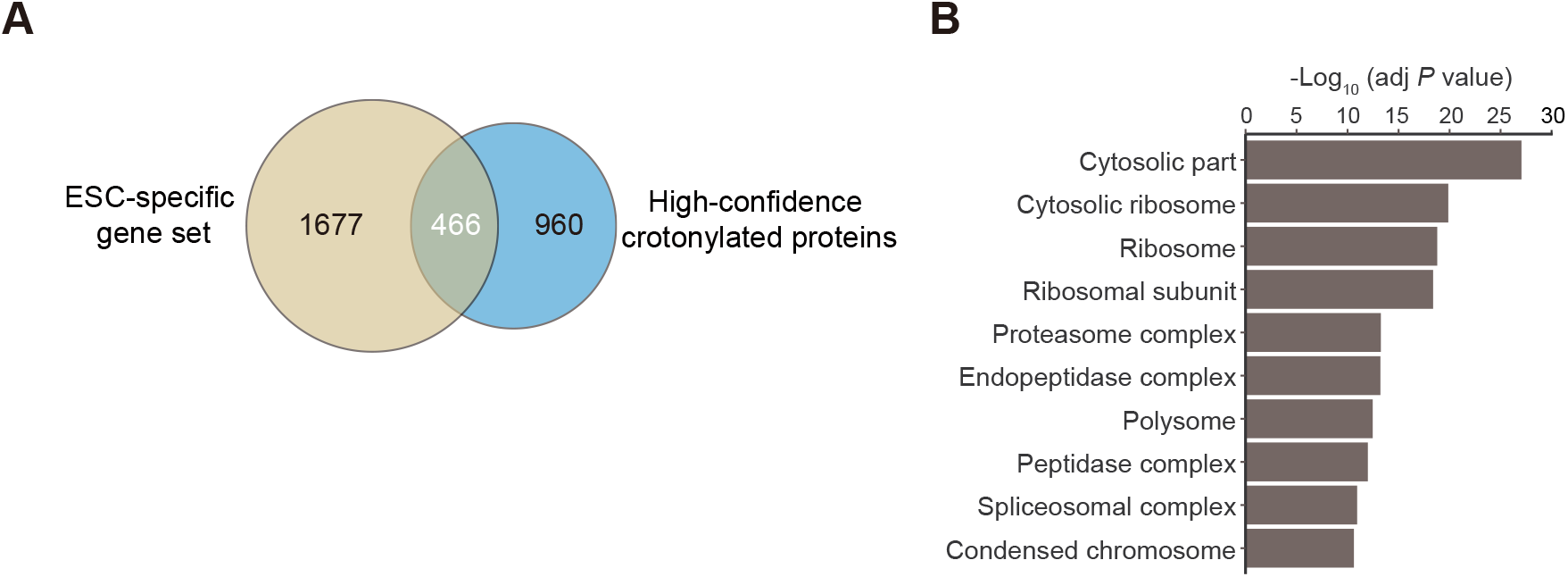
Enrichment for ESC-specific proteins in our crotonylome. *Related to Figure 4*. **A**. Venn diagram comparing our crotonylome datasets with the ‘ESC-specific gene sets’. **B**. GO terms for the 466 overlapping proteins from (**A**) (Fisher’s exact test, Benjamini-Hochberg corrected *P* < 0.01).

